# Whole-Brain Longitudinal Profiling of Serotonergic Reuptake Inhibition

**DOI:** 10.1101/2020.08.10.243899

**Authors:** Horea-Ioan Ioanas, Bechara John Saab, Markus Rudin

## Abstract

The serotonergic system is widely implicated in affect regulation, and a common target for psychopharmacological interventions. Selective Serotonin Reuptake Inhibitors (SSRIs) are the foremost drug class for treating depression, as well as anxiety, phobia and other affective disorders. However, the functional mechanisms determining SSRI efficacy remain elusive, hindering the targeted further development of serotonergic system interventions. Assays for longitudinal whole-brain interrogation of the serotonergic system are unavailable, yet such techniques are essential for identifying differential intervention effects across projection areas. We present a novel longitudinal opto-fMRI assay suitable for imaging longitudinal drug treatment effects on the mouse serotonergic system — within-subject and with sub-millimetre spatial resolution. We apply this assay to a longitudinal fluoxetine treatment, and document reliable segmentation of brain-wide treatment effects, including identification of a brainstem cluster with a highly significant longitudinal trajectory, constituting a novel neurophenotype for psychopharmacological interventions. We differentiate serotonergic neuron activation from projection area activation, and offer brain-wide fMRI evidence for the prominent autoinhibition down-regulation theory of SSRI effects. Further, we show that given the sensitivity of the assay, SSRI treatment produces no persistent effects after treatment cessation in healthy subjects.

## Introduction

The serotonergic system comprises neurons defined by the production and secretion of the neurotransmitter serotonin (5-HT). This evolutionary highly conserved system, with its hub in the nonlateralized brainstem dorsal raphe nucleus (DR), encompasses around 9000 neurons in the mouse [1] and 11500 in the rat [2]. Though the small number of neurons obscures the system in tractography analysis, it projects to a large subset of brain regions (fig. 1a), exerting significant control over them [3], and thus constitutes an important node in the graph representation of the brain.

**Figure 1:**
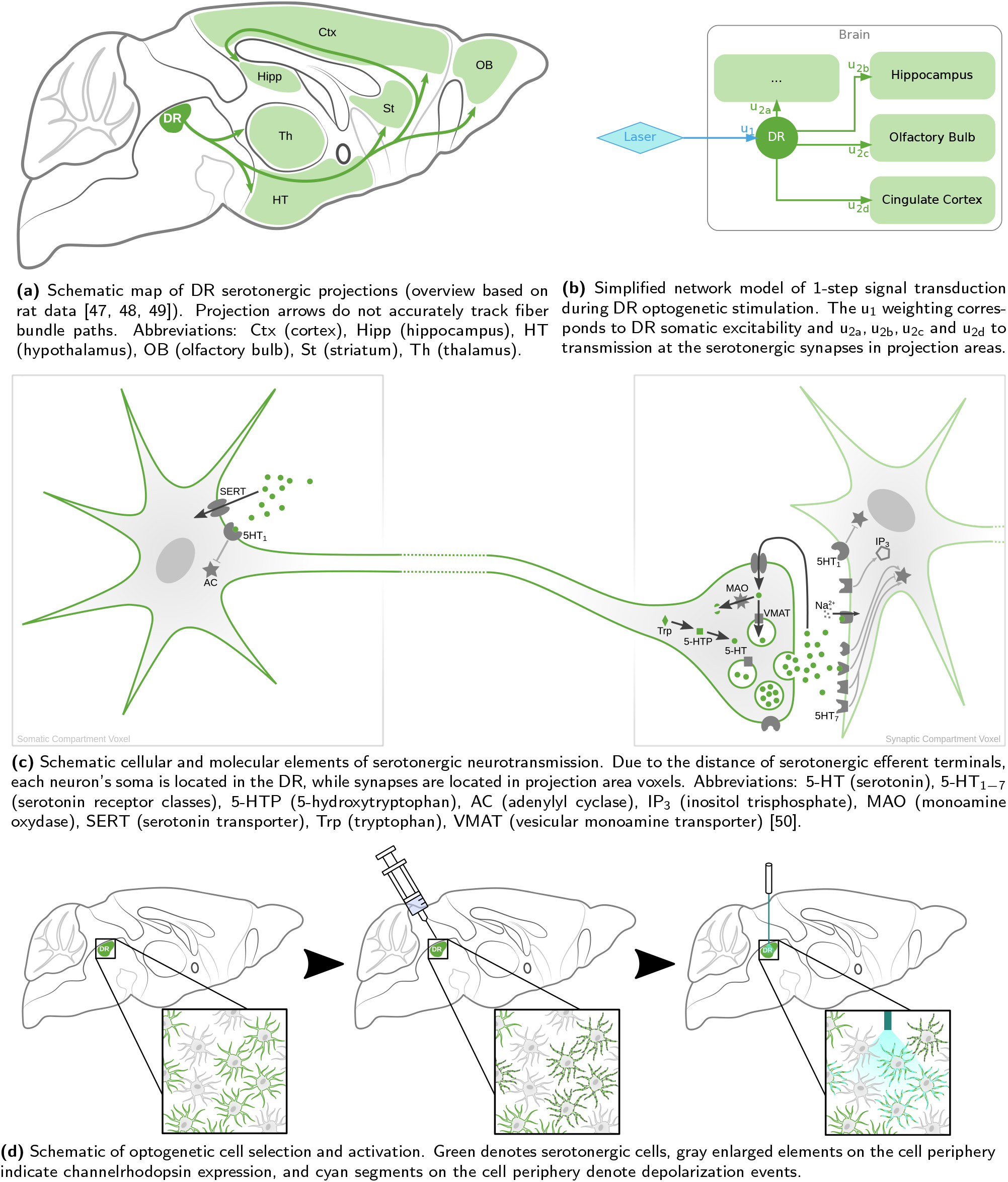
The cellular compartmentalization of serotonergic neurotransmission and potential pharmacological targets can partly be mapped onto neuroanatomical features by a simple network model, using optogenetics. Depicted are schematic overviews of the DR ascending serotonergic system at various spatial resolutions and abstraction levels.

Consistent with wide-ranging projections, the serotonergic system is implicated in numerous cognitive and behavioural phenomena — including impulse control, affect, social behaviour, and in particular social dominance [4, 5]. Since most of these phenomena are not under direct cognitive control, exogenous control of the neuronal systems that underpin them is highly sought after.

Perhaps the most prominent phenomenon in which the serotonergic system is implicated is affect, of which the most common dysfunction is depression. The implication of the serotonergic system in depression is highlighted by serotonin transporter promoter polymorphisms correlating with depression incidence [6]. This is corroborated by the observation that selective serotonin reuptake is implicated in the etiology of depression [7], and that the inhibition thereof is a viable treatment option for depression [8].

In fact, the drugs most commonly used for the treatment of depression are serotonin reuptake inhibitors, such as the selective serotonin reuptake inhibitor (SSRI), fluoxetine [9]. SSRI treatment owes its success to the targeted and homeostasis-modulating manner in which reuptake inhibition influences serotonergic function. This is in opposition to direct targeting via receptor agonists or releasing agents — which can cause extensive non-physiologic side effects [10] on account of eliciting activity at a rate only limited by receptor density and distribution, or cause serotonergic depletion [11], respectively. In line with the homeostatis-modulating interpretation of SSRIs effects, drugs of this class are documented to only produce therapeutic effects after a 1-2 week period [12].

The most prominent theory for serotonergic homeostasis modulation is based on down-regulation of autoinhibition via 5-HT_1_ class receptors [13, 14]. This model states that, during SSRI treatment, autoinhibitors chronically receive atypically high feedback signal upon serotonin release. In response to this signal, autoinhibitors become down-regulated and/or desensitized in their function, thereby increasing the excitability of serotonergic neurons. This theory is supported by functional as well as protein-expression data [15, 16], but there remains a significant gap in understanding how such a mechanism impacts serotonergic signalling at the whole-brain level.

Assessing whole-brain function in vivo is a challenging task, as most measurement techniques suited to interrogate neurotransmitter or cell-type-specific signalling (microdialysis and electrochemistry, or electrophysiology and optical imaging, respectively) either lack spatial resolution entirely, or are significantly limited in depth penetration. Functional magnetic resonance imaging (fMRI), while comparatively unspecific in terms of the cellular processes being imaged, provides both sub-millimeter spatial resolution, as well as full brain coverage without the need for invasive intervention, and is thus well-suited for longitudinal application. While hemodynamic contrast constitutes only an indirect measure of neuronal function, additional specificity can be obtained in animal models by targeting specific neurotransmitter systems via optogenetics. Further, model animal research (as well as translational integration of evidence across model organisms) is particularly suitable for the serotonergic system, owing to its high evolutionary conservation [17].

Serotonergic cell-type specific stimulation in the DR, concurrent with opto-fMRI, can thus be used to highlight neurotransmitter system function at the whole brain level. This assay can be iterated over a longitudinal drug treatment period, and thus verify whether an SSRI (e.g. fluoxetine) induces meaningful changes on the highlighted system. Given such time-dependent effects, it can further be ascertained whether they are representative of the therapeutic profile of SSRIs, and compatible with the autoinhibition down-regulation theory. Given the significant extent to which SSRIs are prescribed, the question of whether and how SSRI treatment affects the function of the healthy brain can also be addressed, by using a healthy model animal population. Of particular relevance in the context of SSRI effects in healthy subjects, is the question of whether SSRI effects persist post treatment in any of the brain regions.

The interpretation of results with regard to molecular mechanisms is rendered particularly convenient by the notable length of the projections (fig. 1a). Given such a spatial distribution, distinct voxels can easily be mapped onto cellular components (fig. 1c), with DR voxels corresponding to neuronal somata, and projection area voxels corresponding to synaptic compartments. Based on such a mapping, a simple network model (fig. 1b) can be used to interpret the results of signal propagation from the initial excitatory stimulation delivered to the DR. In a simple stimulus-evoked analysis, voxel behaviour in the DR thus represents serotonergic cell excitability. By contrast, voxel behaviour in projection areas represents the sum of serotonergic cell excitability and transmission strength at the serotonergic synapse. More unambiguous estimates of transmission can be obtained via seed-based functional connectivity, yet the analysis method may be significantly more susceptible to noise than stimulus-evoked analysis [18].

## Materials and Methods

### Animal Preparation

DR serotonergic neurons were specifically targeted via optogenetic stimulation. As shown in fig. 1d, this process entails a three-stage selection process: the cell-type selection based on the transgenic mouse strain, the location selection based on the injection site, and the activation based on the overlap of the aforementioned selection steps with the optic fiber tip. The study was performed in mice expresses Cre recombinase under a Pet-1 transcription factor enhancer (ePet), which is uniquely expressed in serotonergic neurons [19]. Construct presence was assessed via polymerase chain reaction (PCR) of a Cre gene segment, using the forward primer ACCAGCCAGCTATCAACTCG and the reverse primer TTGCCCCTGTTTCACTATCC.

The DR of the animals was injected with a solution containing recombinant Adeno-Associated Viruses (rAAVs). The injection entry point was 0.6 mm caudal and 1 mm right of lambda, with the cannula being inserted to a depth of 3.6 mm from the skull at a yaw of 20° towards the left of the dorsoventral axis. The vector delivered a plasmid containing a floxed channelrhodopsin and YFP construct: pAAV-EF1a-double floxed-hChR2(H134R)-EYFP-WPRE-HGHpA, gifted to a public repository by Karl Deisseroth (Addgene plasmid #20298). Viral vectors and plasmids were produced by the Viral Vector Facility (VVF) of the Neuroscience Center Zurich (Zentrum für Neurowissenschaften Zürich, ZNZ). The solution was prepared at a titer of 5.7 × 10^12^ vg/ml and the injection volume was 1.3 μl. Construct expression was ascertained post mortem via fluorescence microscopy of formaldehyde fixated 200 μm brain slices.

Animals were fitted with an optic fiber implant (l = 3.3 mm, d = 400 μm, NA = 0.22) inserted perpendicularly, to its full length, at 0.6 mm caudal of lambda and on the midline.

All animal work was performed in accordance with relevant animal welfare regulations as implemented by the Cantonal Veterinary Office of Zurich.

### Drug Treatments

The effects of acute and chronic fluoxetine treatment were evaluated based both on novel drinking water administration data published herein, and on previously published (but, as of yet, unevaluated) intraperitoneal administration data [20]. A graphical overview for the time courses of both these treatments can be found in fig. S1. For both chronic treatment experiments, fluoxetine hydrochloride (≥99 % HPLC, Bio-Techne AG, Switzerland) was used to prepare an injection solution (2.25 mg/ml in saline). For the acute fluoxetine administration session of both treatment experiments, a volume of the solution adjusted to deliver fluoxetine at 10 mg/(kg BW) was injected intravenously 10 min prior to functional scan acquisition.

#### Chronic Drinking Water Administration Experiment

Fluoxetine hydrochloride (≥99 % HPLC, Bio-Techne AG, Switzerland) was used to prepare a drinking water (124 mg/l in tap water) solution. Drinking behaviour was monitored to ascertain daily fluoxetine consumption of not fewer than 15 mg/(kg BW) and the intravenous solution volume was adjusted to deliver fluoxetine at 10 mg/(kg BW). Drinking water during and prior to the experiment was delivered from bottles wrapped in aluminium foil. The drinking solution as well as the fodder, cage, and bedding were replaced weekly — in synchrony with the measurements (after each measurement, and at corresponding intervals between measurements). This dataset includes fMRI recordings from 15 animals subjected to fluoxetine treatment and 14 animals serving as vehicle administration controls.

#### Chronic Intraperitoneal Administration Experiment

Over the course of the chronic treatment period (fig. S1b), the fluoxetine injection solution was delivered intraperitoneally once a day, every day. The volume for chronic (intraperitoneal) injections was adjusted to deliver fluoxetine at 10 mg/(kg BW). This dataset includes fMRI recordings from 11 animals, all subjected to fluoxetine treatment.

### MR Acquisition

Over the course of preparation and measurement, animals were provided a constant flow of air with an additional 20 % O_2_ gas (yielding a total O_2_ concentration of ≈36 %). For animal preparation, anesthesia was induced with 3 % isoflurane, and maintained at 2 to 3 % during preparation — contingent on animal reflexes. Animals were fixed to a heated MRI-compatible cradle via ear bars and a face mask equipped with a bite hook. A subcutaneous (s.c.; right dorsal) and intravenous (i.v.; tail vein) infusion line were applied. After animal fixation, a bolus of medetomidine hydrochloride (Domitor, Pfizer Pharmaceuticals, UK) was delivered s.c. to a total dose of 100 ng/(g BW) and the inhalation anesthetic was reduced to 1.5 % isoflurane. After a 5 min interval, the inhalation anesthetic was set to 0.5 % and medetomidine was continuously delivered at 200 ng/(g BW h) for the duration of the experiment. This anesthetic protocol is closely based on extensive research into animal preparation for fMRI [21].

All data were acquired with a Bruker Biospec system (7 T, 16 cm bore), and an in-house built transmit/receive surface coil, engineered to permit optic fiber implant protrusion.

#### Chronic Drinking Water Administration Experiment

Anatomical scans were acquired via a TurboRARE sequence, with a RARE factor of 8, an echo time (TE) of 30 ms, an inter-echo spacing of 10 ms, and a repetition time (TR) of 2.95 s. Thirty adjacent (no slice gap) coronal slices were recorded with a nominal in-plane resolution of Δx(*ν*) = Δy(*ϕ*) = 75 μm, and a slice thickness of Δz(t) = 450 μm.

Functional scans were acquired with a gradient-echo EPI sequence, a flip angle of 60°, and TR/TE = 1000 ms/5.9 ms. Thirty adjacent (no slice gap) coronal slices were recorded with a nominal in-plane resolution of Δx(*ν*) = Δy(*ϕ*) = 225 μm, and a slice thickness of Δz(t) = 450 μm. Changes in cerebral blood volume (CBV) are measured as a proxy of neuronal activity following the administration of an intravascular iron oxide nanoparticle based contrast agent (Endorem, Laboratoire Guebet SA, France). The contrast agent (30.24 μg/(g BW)) was delivered as an i.v. bolus 10 min prior to the fMRI data acquisition, to achieve a pseudo steady-state blood concentration. This contrast is chosen to enable short echo-time imaging thereby minimizing artefacts caused by gradients in magnetic susceptibility [22].

The DR was stimulated via an Omicron LuxX 488-60 laser (488 nm) tuned to a power of 30 mW at contact with the fiber implant, according to the protocol listed in table S1.

#### Chronic Intraperitoneal Administration Experiment

The acquisition parameters of this dataset are detailed in the original publication [20]. The DR was stimulated via an Omicron LuxX 488-60 laser (488 nm) tuned to a power of 30 mW at contact with the fiber implant, according to the protocol listed in table S2.

### Data Processing

Stimulation protocols were delivered to the laser and recorded to disk via the COSplayer device [23]. Animal physiology, preparation, and measurement metadata were tracked with the LabbookDB database framework [24].

Data conversion from the proprietary ParaVision format was performed via the Bruker-to-BIDS repositing pipeline [25] of the SAMRI package (version 0.4 [26]). Following conversion, data was dummy-scan corrected, registered, and subject to controlled smoothing via the SAMRI Generic registration workflow [20]. As part of this processing, the first 10 volumes were discarded (automatically accounting for volumes excluded by the scanner software). Registration was performed using the standard SAMRI mouse-brain-optimized parameter set for ANTs [27] (version 2.3.1). Data was transformed to a stereotactically oriented standard space (dsurquec, as distributed in the Mouse Brain Atlases Package [28], version 0.5.3), which is based on a high-resolution T_2_-weighted atlas [29]. Controlled spatial smoothing was applied in the coronal plane up to 250 μm via the AFNI package [30] (version 19.3.12).

The registered time course data was frequency filtered depending on the analysis workflow. For stimulus-evoked activity, the data was low-pass filtered at a period threshold of 225 s, and for seed-based functional connectivity, the data was band-pass filtered within a period range of 2 to 225 s.

### Statistics

Volumetric data was modelled using functions from the FSL software package [31] (version 5.0.11). First-level regression was applied to the temporally resolved volumetric data via FSL’s glm function, whereas the second-level analysis was applied to the first-level contrast and variance estimates via FSL’s flameo.

Stimulus-evoked first-level regression was performed using a convolution of the stimulus sequence with an opto-fMRI impulse response function, estimated by a beta fit of previously reported mouse opto-fMRI responses [3]. Seed-based functional connectivity analysis was performed by regressing the time course of the DR voxel most sensitive to stimulus-evoked activity (per scan).

Brain parcellation for region-based evaluation was performed using a non-overlapping multi-center labelling [29, 32, 33, 34], as distributed in version 0.5.3 of the Mouse Brain Atlases data package [28]. The mapping operations were performed by a SAMRI function, using the nibabel [35] and nilearn [36] libraries (versions 2.3.1 and 0.5.0, respectively). Gaussian mixture modelling for voxel classification was performed via the GaussianMixture class from the scikit-learn library [37] (version 0.20.3). Distribution density visualizations were created using the Scott band-width density estimator [38]. For the visualization of parcellation-based statistic score density distributions, regions with a volume lower than 0.06 mm^3^ or with more than 34 % exact zero values are excluded from analysis.

Higher-level statistical modelling was performed with the Statsmodels software package [39] (version 0.9.9), and the SciPy software package [40] (version 1.1.0). Model parameters were estimated using the ordinary least squares method, and a type 3 analysis of variance (ANOVA) was employed to control estimate variability for unbalanced categories. All t-tests producing explicitly noted p-values are two-tailed, and all post-hoc t-tests use the Benjamini-Hochberg procedure, controlling the false discovery rate (FDR) at *α* = 0.05.

Software management relevant for the exact reproduction of the aforementioned environment was performed via neuroscience package install instructions for the Gentoo Linux distribution [41].

### Reproducibility and Open Data

The resulting t-statistic maps, as well as the GMM cluster assignment maps (both selected and rejected) are distributed along the source-code of all analyses [42]. The BIDS [43] data archives which serve as raw data recourse for this document are openly distributed [44, 45], as is the full instruction set for recreating this document [42]. The source code for this document and all data analysis shown herein is published according to the RepSeP specifications [46]. The data analysis execution and document compilation has been tested repeatedly on numerous machines, and as such we attest that the figures and statistics presented can be reproduced based solely on the raw data, dependency list, and analysis scripts which we distribute.

## Results

The activation pattern elicited by photic stimulation of transfected DR neurons shows strong inhibition of cortical areas (fig. 2c) and activation of subcortical areas, primarily in the brainstem (fig. 2d). Notably, due to extended brain coverage compared to past studies, we also resolve cerebellar regions, where we also observe activation. The most salient activation peak is seen ventral of the sylvian aqueduct, in the DR target area, and features a concavity at the dorsal delimitation, approximately at the position of the optic implant.

**Figure 2:**
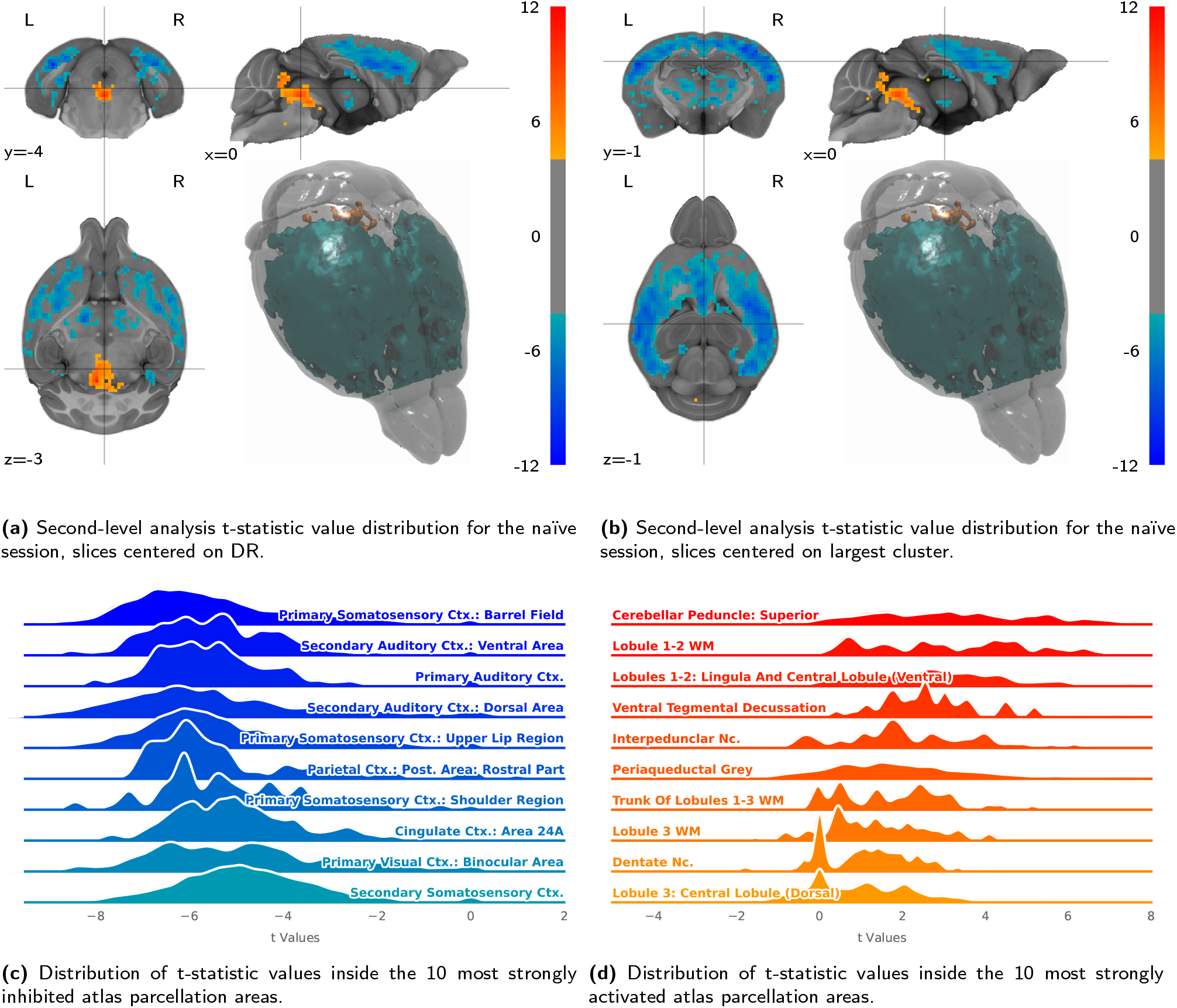
DR stimulation in the drug-naïve condition elicits a large inhibition cluster in cortical regions and focused activation in and around the DR. Presented are baseline activation maps **(a, b)**, evaluated according to atlas parcellation regions **(c, d)**. Abbreviations: Ctx. (Cortex), Nc. (Nucleus), WM (White Matter).

We observe no significant laterality main effect or laterality-session interaction in mice exposed either to drinking water (*F*_1,2852_ = 0.091, *p* = 0.76, *F*_4,2852_ = 0.074, *p* = 0.99) or intraperitoneal (*F*_1,1416_ = 0.42, *p* = 0.52, *F*_4,1416_ = 0.11, *p* = 0.98) fluoxetine administration. This observation warrants a hemisphere-agnostic atlas parcellation for both of these two datasets.

Longitudinal analysis of the DR target stimulation site (fig. 5a) in the drinking water administration dataset reveals no significant main effect for the session (*F*_1,92_ = 0.27, *p* = 0.61), but a significant categorical session-treatment interaction effect (*F*_9,92_ = 6.365, *p* = 5.65 × 10^−7^). Post-hoc t-tests reveal that the interaction effect is significant at 2 weeeks (*p* = 9×10^−3^) and 4 weeks (*p* = 4×10^−3^) of chronic administration, but not the for the naïve (*p* = 0.79), acute (*p* = 0.15), or post-treatment (*p* = 0.77) sessions. An equivalent longitudinal pattern is not seen in the intraperitoneal administration data (fig. S3a).

To identify coherent longitudinal trajectory clusters outside of the primary DR target area, we apply unsupervised Gaussian mixture modelling (GMM) to a 5-dimensional representation of session-wise second-level maps for the fluoxetine treatment group. As cross-validation is ill-defined for unsupervised learning methods, we use classification reliability estimation (fig. 3a) for model selection. We set up a model pool exploring up to 15 clusters and all available covariance structures. We observe the highest classification reliabilities for 4-cluster and 5-cluster models using a spherical covariance structure. Additionally, we observe an increase in classification reliability for 9 to 12 components in models using a tied covariance structure — which indicates strong inter-session correlation in smaller clusters.

**Figure 3:**
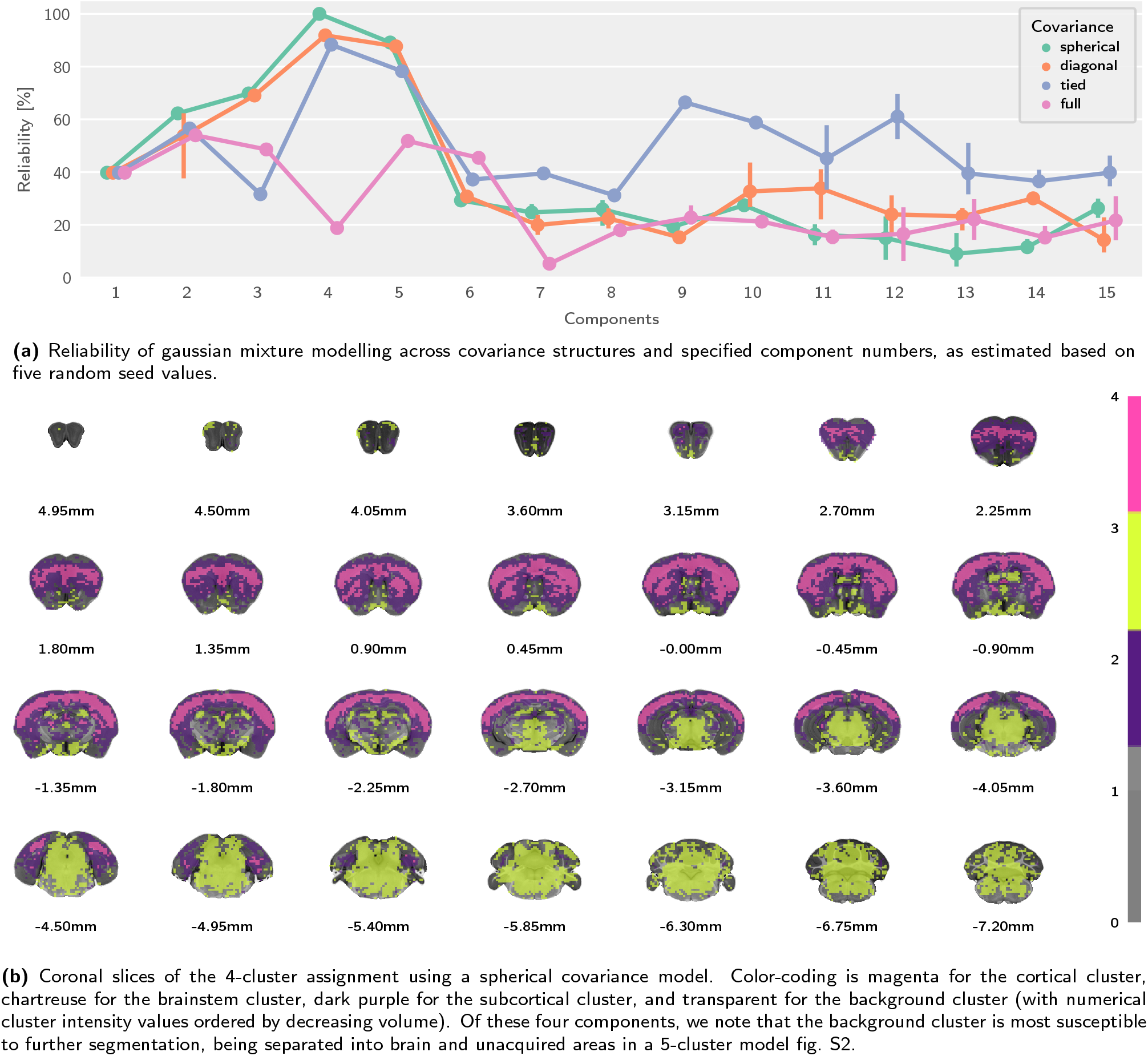
Gaussian mixture modelling reveals highly stable voxel assignment into four or five clusters. Depicted is the reliability of the model variants **(a)** as well as a volumetric plot of the most reliable classification **(b)**.

The selected clustering model (fig. 3b, using 4 clusters and spherical covariance) identifies a *background cluster* (which includes rostral and caudal brain areas, not covered by data acquisition), a *cortical cluster*, a *brainstem cluster*, and a *subcortical cluster*. To estimate the significance of the longitudinal trajectories for each cluster, we apply them as masks to both the fluoxetine and vehicle treatment data and model the cluster-wise means from first-level subject-wise general linear modelling (GLM) results.

The cortical cluster extends over the majority of cortical areas, also curving downward along the midline into cingulate cortex areas (fig. 4a). The cortical parcellation areas with the highest proportion of cortical cluster voxels belong to the somatosensory and cingulate cortex (fig. 4b). Longitudinal analysis of the cluster region of interest (fig. 5b) in the drinking water administration dataset reveals no significant main effect for the session (*F*_1,92_ = 0.28, *p* = 0.6), but a significant categorical session-treatment interaction effect (*F*_9,92_ = 11.2, *p* = 1.22 × 10^−11^). Post-hoc t-tests reveal that the interaction effect is significant at the acute administration (*p* = 1×10^−4^) and 2 weeek chronic administration (*p* = 0.033) levels, but not the 4 week (*p* = 0.12) chronic administration level, nor the the naïve (*p* = 0.33) or post-administration (*p* = 0.69) levels. This longitudinal pattern is also discernible in the intraperitoneal administration data (fig. S3b) — though the trajectory more closely resembles a biphasic (acute vs. chronic) response.

**Figure 4:**
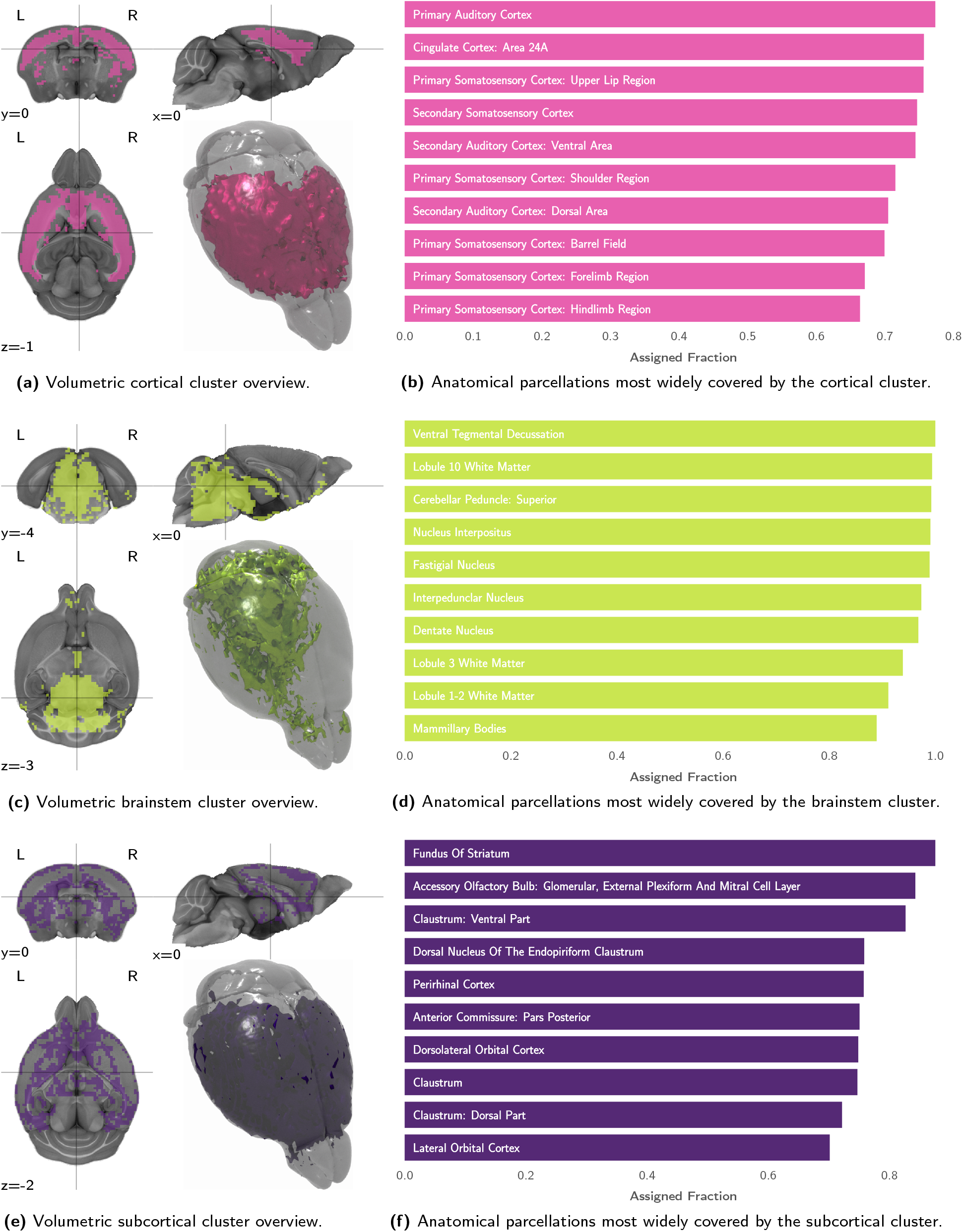
The longitudinal trajectory clusters show a spatial distribution following a brainstem-cortex separation. Depicted are volumetric representations sliced on the respective cluster center of mass **(a, c, e)**, and anatomical parcellation overlap proportions **(b, d, f)**.

**Figure 5:**
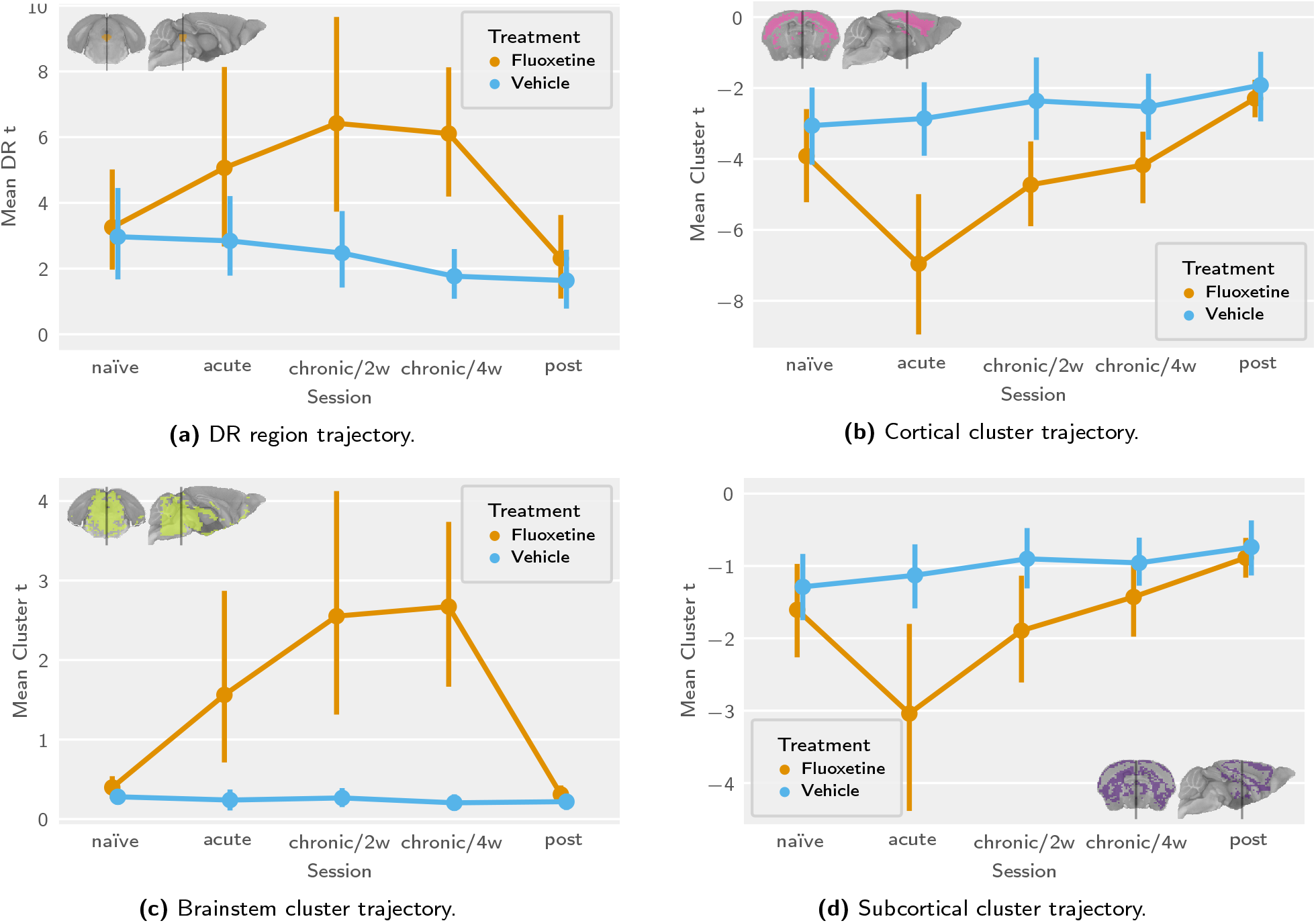
Longitudinal trajectories primarily show chronic SSRI sensitivity for the DR and brainstem clusters, and acute SSRI sensitivity for the cortical and subcortical clusters. Depicted are activation time courses in the drinking water administration dataset, for highlighted regions, including the DR **(a)** and longitudinal trajectory clusters **(b, c, d)**.

The brainstem cluster extends caudally into the cerebellum, and rostrally into the thalamus, hypothalamus, and along the ventral aspect of the brain (fig. 4c). The cluster fully or almost fully covers numerous atlas parcellation areas of the brainstem and rostral cerebellum (fig. 4d). Longitudinal analysis of the cluster region of interest (fig. 5c) in the drinking water administration dataset reveals a significant main effect for the session (*F*_1,92_ = 20.67, *p* = 1.65 × 10^−5^) and a significant categorical session-treatment interaction effect (*F*_9,92_ = 9.533, *p* = 3.90 × 10^−10^). Post-hoc t-tests reveal that the interaction effect is significant at the acute administration level (*p* = 0.012), as well as the 2 week (*p* = 3×10^−5^) and 4 week (*p* = 5×10^−6^) chronic administration levels — but not the naïve (*p* = 0.86), or post-administration (*p* = 0.86) levels. This longitudinal pattern is not well-discernible in the intraperitoneal administration data (fig. S3c).

The subcortical cluster brackets the cortex (as a thin shell, covering the deepest and most superficial regions of the cortex). It extends into rostral subcortical areas, roughly following anatomical features, such as the globus pallidus (fig. 4e). The cluster does not fully encompass many anatomical parcellation areas, but of the areas into which it extends, striatal areas are the most prominent (fig. 4f). Longitudinal analysis of the cluster region of interest (fig. 5d) in the drinking water administration dataset reveals no significant main effect for the session (*F*_1,92_ = 0.87, *p* = 0.35), but a significant categorical session-treatment interaction effect (*F*_9,92_ = 7.196, *p* = 7.62 × 10^−8^). Post-hoc ests reveal that the interaction effect is significant at the acute administration level (*p* = 6×10^−4^), but not the naïve (*p* = 0.54), 2 weeek (*p* = 0.11) and 4 week (*p* = 0.53) chronic administration, or post-administration (*p* = 0.76) level. This longitudinal pattern is also discernible in the intraperitoneal administration data (fig. S3d), again with a trajectory resembling a biphasic (acute vs. chronic) response.

## Discussion

In this article we interrogate the serotonergic system in the mouse over a longitudinal course of fluoxetine treatment, by using optogenetic stimulation of serotonergic DR neurons in conjunction with fMRI. Activation patterns observed at baseline (figs. 2a, 2c and 2d) were analogous to those reported in a previous study [3]. We observe that the system shows no lateral preference in its functional effects, neither at baseline, nor during fluoxetine treatment or after treatment cessation.

Our results suggest that fluoxetine treatment induces functional serotonergic effects corroborating the autoinhibition down-regulation theory. This is best represented by the differential temporal trajectories of the DR (fig. 5a) and the cortical projection areas (fig. 5b). In this comparison, the DR becomes significantly more sensitive to stimulation in the chronic, but not the acute administration sessions. In contrast, the cortical projection areas show significantly larger negative signal amplitudes for the acute and 2-week chronic, but not the 4-week chronic sessions.

In order to identify coherent trajectories at the whole brain level we use unsupervised machine learning to determine an appropriate segmentation of voxels into clusters, based on their temporal profile. Acknowledging the limitations inherent in unsupervised model selection, we inspect classification reliability and further provide the cluster assignment maps for all tested models in volumetric form for download [42]. The clusters identified using the most reliable classification parameters show strong spatial coherence, and roughly follow anatomical features (fig. 3b). We name these trajectory-based clusters “brainstem”, “sub-cortical”, and “cortical”.

The cortical trajectory cluster encompasses primarily cortical regions (figs. 4a and 4b), with its temporal profile showing a significant fluoxetine effect upon acute fluoxetine administration, and again in the 2-week chronic administration session. All effects manifest themselves as enhanced inhibition, with the acute effect being notably the highest in amplitude. This is consistent with homeostatic adaptation of the serotonergic system, in the context of which acute exposure to the drug leads to accumulation of neurotransmitter at the synapse and strengthens the inhibitory effect — whereas during chronic administration the synapse adjusts to elicit a postsynaptic effect more consistent with the baseline. Whether this homeostatic effect converges on the baseline or an intermediary stable state (as suggested by the intraperitoneal administration dataset, fig. 5b) is unclear given the duration of chronic administration.

The brainstem trajectory cluster encompasses both the DR and the majority of the rest of the brainstem, the part of the cerebellum captured during data acquisition, some of the ventralmost regions of the forebrain along the midline (fig. 4c), and notably, the fimbria (fig. 3b). The trajectory of this cluster strongly resembles that of the DR, though at a roughly 50 % reduced amplitude (fig. S3c). Interestingly, in this case, significant fluoxetine effects are also found for the acute session. These results are not strictly consistent with the autoinhibition down-regulation theory, but also not necessarily at odds with it. The existence of such a trajectory cluster (including but also extending far beyond the DR) indicates that there are numerous brain areas other than the DR, in which the same sensitization effect can be observed. One possible explanation is that these areas do not experience homeostatic adaptation to increased serotonergic signalling at the synapse, and consequently mirror the same signal behaviour as their afferents. Interestingly, this is the only cluster showing positive signal transmission, and does so only during the treatment window. It can thus be speculated that excitatory serotonergic synapses experience a distinctly evolution during fluoxetine treatment.

The subcortical trajectory cluster is the least spatially coherent, but shows some anatomical alignment with striatal and limbic regions (figs. 4e and 4f). Though it is the only cluster to extend into the hippocampus proper (the brainstem cluster distinctly covers the fimbria), coverage is sparse, predominantly restricted to the granular layer (fig. 3b). Overall, the temporal trajectory is reminiscent of that shown by the cortical cluster, albeit with lower amplitudes, and only showing a significant effect in the acute administration session. The same homeostatic mechanism as for the cortical cluster can be proposed here. Considering its looser spatial arrangement, however, an alternative explanation can be put forward. Given the strong spatial autocorrelation of fMRI signals, it is possible that this cluster simply captures the intermediary effect between the cortical cluster and the background.

As expected, neither the DR region of interest, nor the trajectory clusters show a significant treatment group contrast for the naïve session. This and the stability of activity across regions of interest in the control condition strongly support the robustness of the assay and reliability of the effects observed. While the assay is able to discern fluoxetine treatment effects with region-specific temporal trajectories, neither the DR nor the trajectory clusters show a significant treatment contrast two weeks after fluoxetine withdrawal fig. S4. We thus conclude that in healthy mice there is no post-treatment fluoxetine effect of similar nature or amplitude to either the acute or chronic treatment effects.

In comparing the drinking water and intraperitoneal chronic administration experiments, we note that only the trajectories of the cortical and subcortical clusters are reproducible. Noting the more limited resolution, more sparse slice positioning, and reduced rostrocaudal coverage, we attribute this in part to the incomplete capture of the DR and brainstem activity profiles. Consequently, we recommend high rostrocaudal coverage with dense slice positioning for robust longitudinal opto-fMRI. Such acquisition improves statistic reliability both by eliminating slice gaps, and by improving co-registration stability. A decreased stability of fluoxetine treatment effects under intraperitoneal chronic administration cannot be ruled out, though it seems less plausible that drug administration procedure effects (e.g. animal stress) would specifically degrade the statistical estimates of the brainstem trajectory.

In this study, seed-based analysis is explored as a modelling alternative for improved differentiation of serotonergic cell excitability and serotonergic synapse transmission efficiency. Given variable levels of activation at the primary stimulation site, increased signal in projection areas cannot unambiguously be attributed to more efficient transduction at the synapse. The theoretical expectation is that projection area signal in a seed-based evaluation would be blunted for sessions in which the stimulation site activation is strongest (i.e. the 2-week and 4-week treatment sessions, fig. 5a). However, seed-based analysis of the data at hand shows an opposite effect, whereby it is primarily the acute treatment session for which the projection area response is blunted (comparing fig. S6e and fig. 5b). Overall, seed based connectivity shows reduced statistical estimates fig. S5 and less distinct longitudinal trajectories (fig. S6). This may indicate that the inability of seed-based connectivity to deliver meaningful disambiguation of excitability and transmission in this study is simply an issue of susceptibility to noise in the stimulation area regressor. We conclude that resolving network dynamics is an important endeavour for stimulus-evoked fMRI— yet may require additional methodological improvements in order to afford insight in excess of stimulus-evoked response modelling.

### Summary

We have applied an emerging opto-fMRI assay to inspect longitudinal changes in serotonergic function elicited by acute and chronic treatment with the SSRI fluoxetine. The analysis revealed three trajectory clusters that follow two different patterns consistent with the autoinhibition down-regulation theory, and further clarifies the spatial distribution of fluoxetine treatment effects. We have shown that the most salient of these trajectories can be observed in two distinct datasets with numerous parameter variations (including drug delivery and MRI acquisition). In particular, we have demonstrated that while treatment effects are highly significant, no persistent changes are seen after treatment cessation in healthy mice. This provides evidence against the proposition that fluoxetine induces stable homeostatic shifts persisting beyond the treatment period. Such a statement can speak both for and against fluoxetine administration (and perhaps SSRI administration in general), depending on whether an intervention time window or a persistent shift in function is the desired result.

We suggest that opto-fMRI based analysis of serotonergic signaling constitutes an attractive assay for profiling serotonergic drugs based on spatio-temporally resolved, whole-brain neurotropic effects. The high level of differentiation between effects induced by fluoxetine, a representative of the SSRI drug class, as compared to the vehicle illustrates the potential of the approach. However, it remains to be determined whether the method provides the sensitivity required to robustly distinguish the responses elicited by different serotonergic drugs recommended for chronic administration (such as different SSRIs).

In the interest of transparency and technology access we provide both the code required to perform all the analyses in this article — as well as the resulting summaries and cluster classifications — for public access [42].

## Author Contributions

HII performed the experiments, data analysis, and drafted the article. BJS drafted the experimental protocols, provided materials and methods, consulted on psychopharmacological methods, and reviewed the article. MR supervised the project, provided materials and methods, consulted on MRI methods, and reviewed and edited the article.

## Supplementary Materials

**Table S1:** Phasic stimulation protocol, coded “CogB”, used in the drinking water administration data.

| Onset<br>[s] | Duration<br>[s] | Frequency<br>[Hz] | Pulse Width<br>[s] |
| --- | --- | --- | --- |
| 182.0 | 20.0 | 20.0 | 0.005 |
| 332.0 | 20.0 | 20.0 | 0.005 |
| 482.0 | 20.0 | 20.0 | 0.005 |
| 632.0 | 20.0 | 20.0 | 0.005 |
| 782.0 | 20.0 | 20.0 | 0.005 |
| 932.0 | 20.0 | 20.0 | 0.005 |
| 1082.0 | 20.0 | 20.0 | 0.005 |
| 1232.0 | 20.0 | 20.0 | 0.005 |

**Table S2:** Phasic stimulation protocol [3], coded “JogB”, used in the intraperitoneal administration data [20].

| Onset<br>[s] | Duration<br>[s] | Frequency<br>[Hz] | Pulse Width<br>[s] |
| --- | --- | --- | --- |
| 222.0 | 20.0 | 20.0 | 0.005 |
| 402.0 | 20.0 | 20.0 | 0.005 |
| 582.0 | 20.0 | 20.0 | 0.005 |
| 762.0 | 20.0 | 20.0 | 0.005 |
| 942.0 | 20.0 | 20.0 | 0.005 |
| 1122.0 | 20.0 | 20.0 | 0.005 |

**Figure S1:**
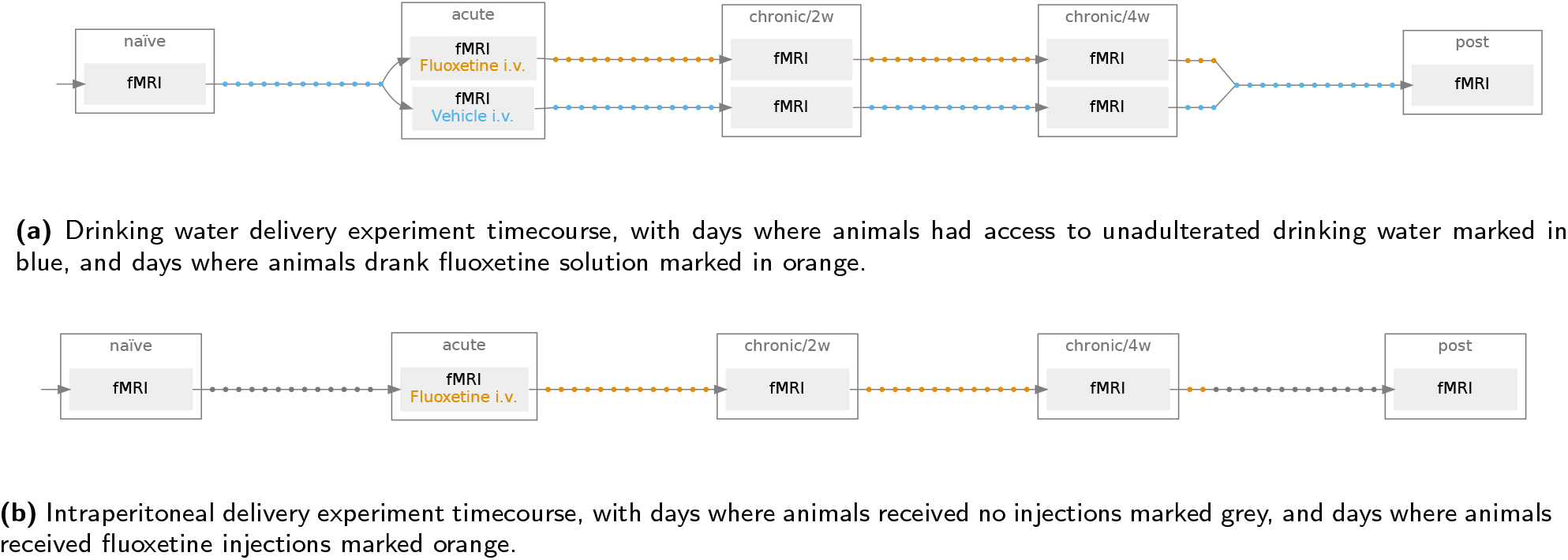
Schematic of the longitudinal experiment timetable design. Opto-fMRI measurement days are denoted via box nodes, and acute drug delivery for the experiment is noted inside the boxes. Grey outer cassettes denote the session identifier names for the study.

**Figure S2:**
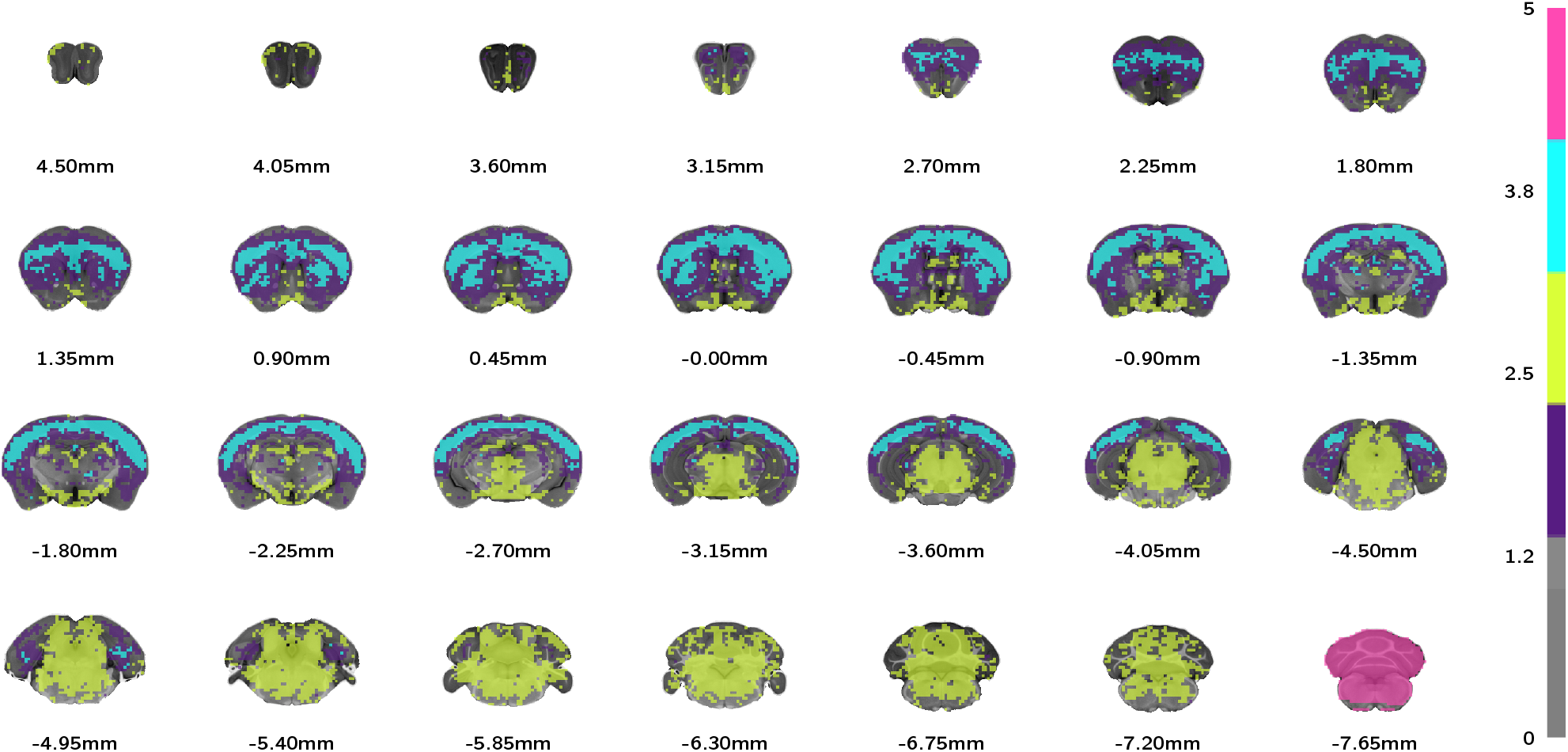
The least stable of the clusters in the 4-cluster model (fig. 3b) is the background cluster. Depicted are coronal slices of the 5-cluster assignment, using a spherical covariance model, showing that the background cluster is further broken down into a brain-tissue background and padded-value (unacquired) region, the latter of which is highlighted in magenta on the last slice.

**Figure S3:**
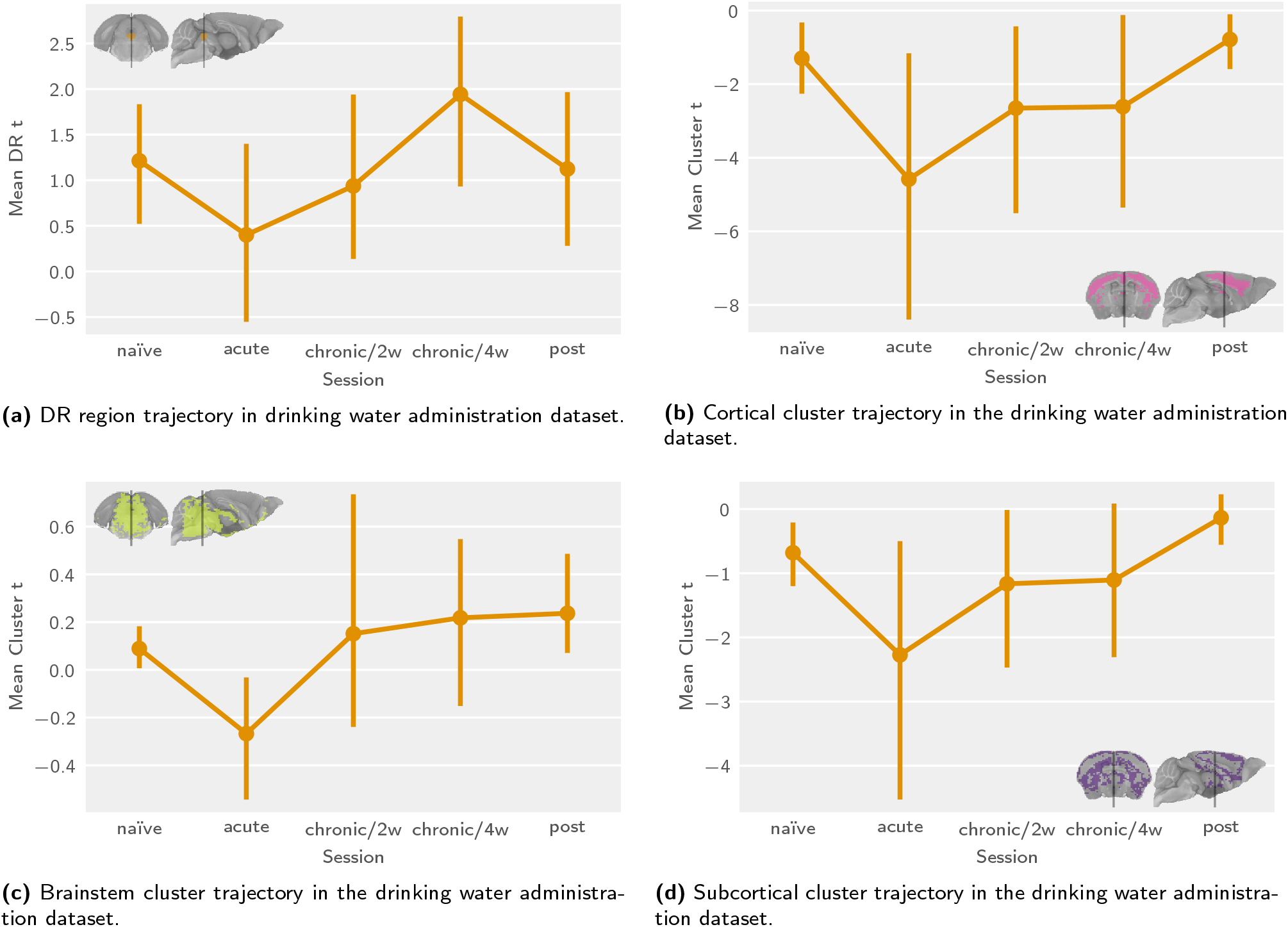
The intraperitoneal administration dataset only detects longitudinal trajectory trends resembling those seen in the drinking water administration dataset for the cortical and subcortical clusters. Depicted are activation time courses for highlighted regions, including the DR **(a)** and longitudinal trajectory clusters **(b, c, d)**.

**Figure S4:**
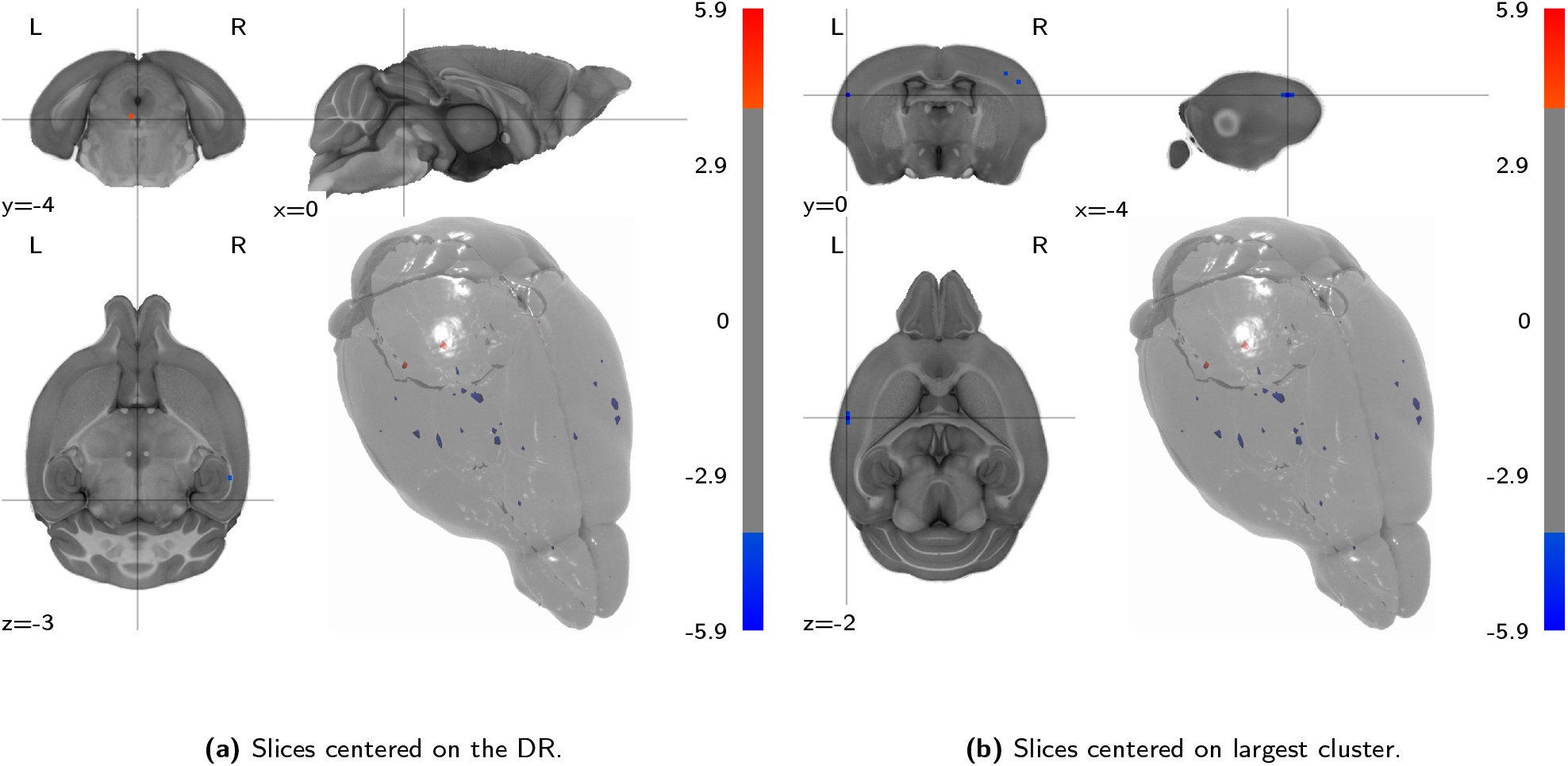
Fluoxetine treatment induces no extensive post-treatment change in stimulus-evoked serotonergic activity. Presented are second-level analysis t-statistic maps for the contrast between the fluoxetine and vehicle groups in the post-treatment session.

**Figure S5:**
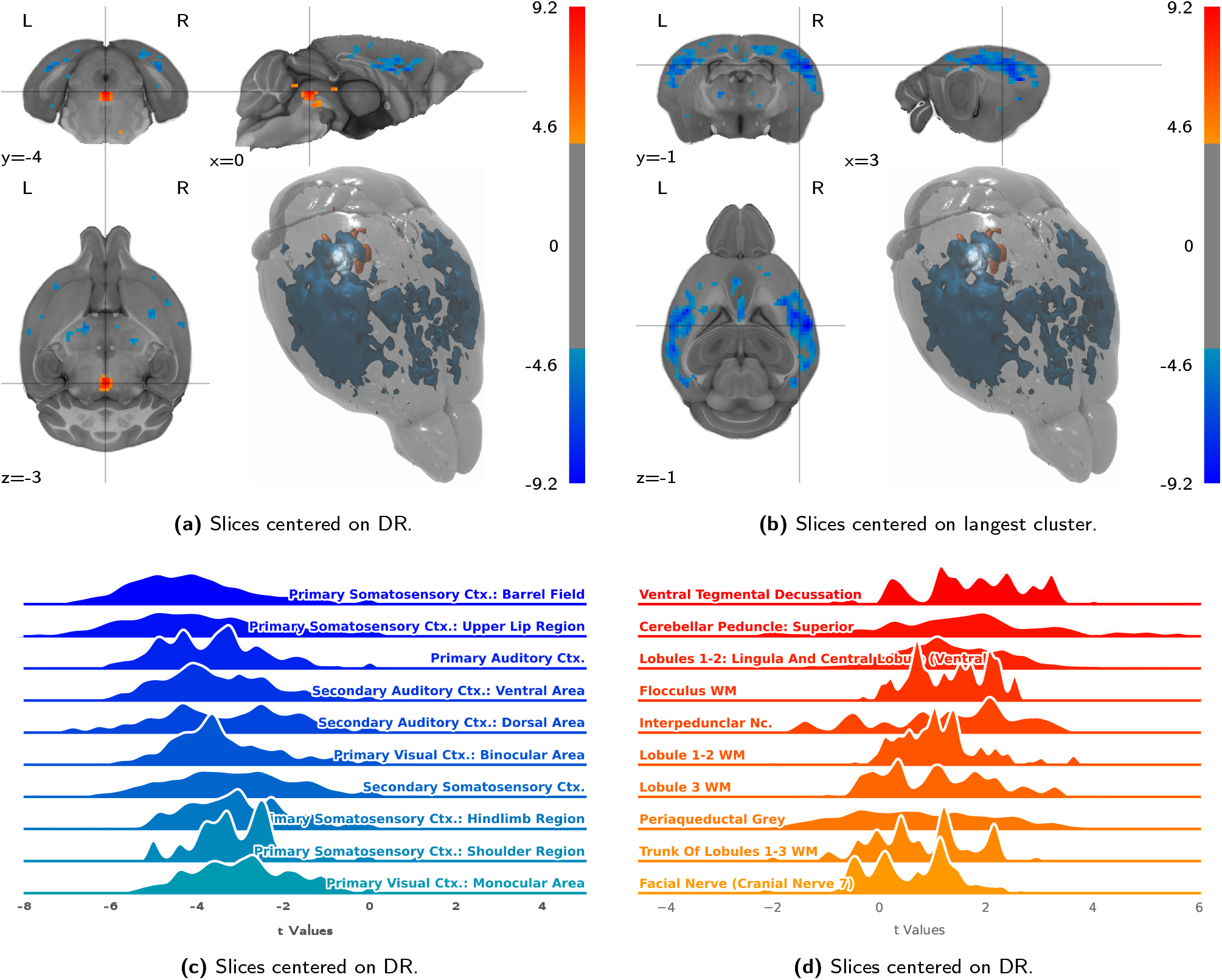
The seed-based connectivity t-statistic pattern closely matches the stimulus-evoked activation pattern, with lower signal intensity. Presented are transmission maps **(a, b)** broken up by atlas parcellation regions **(c, d)**. Abbreviations: Ctx. (Cortex), Nc. (Nucleus), WM (White Matter).

**Figure S6:**
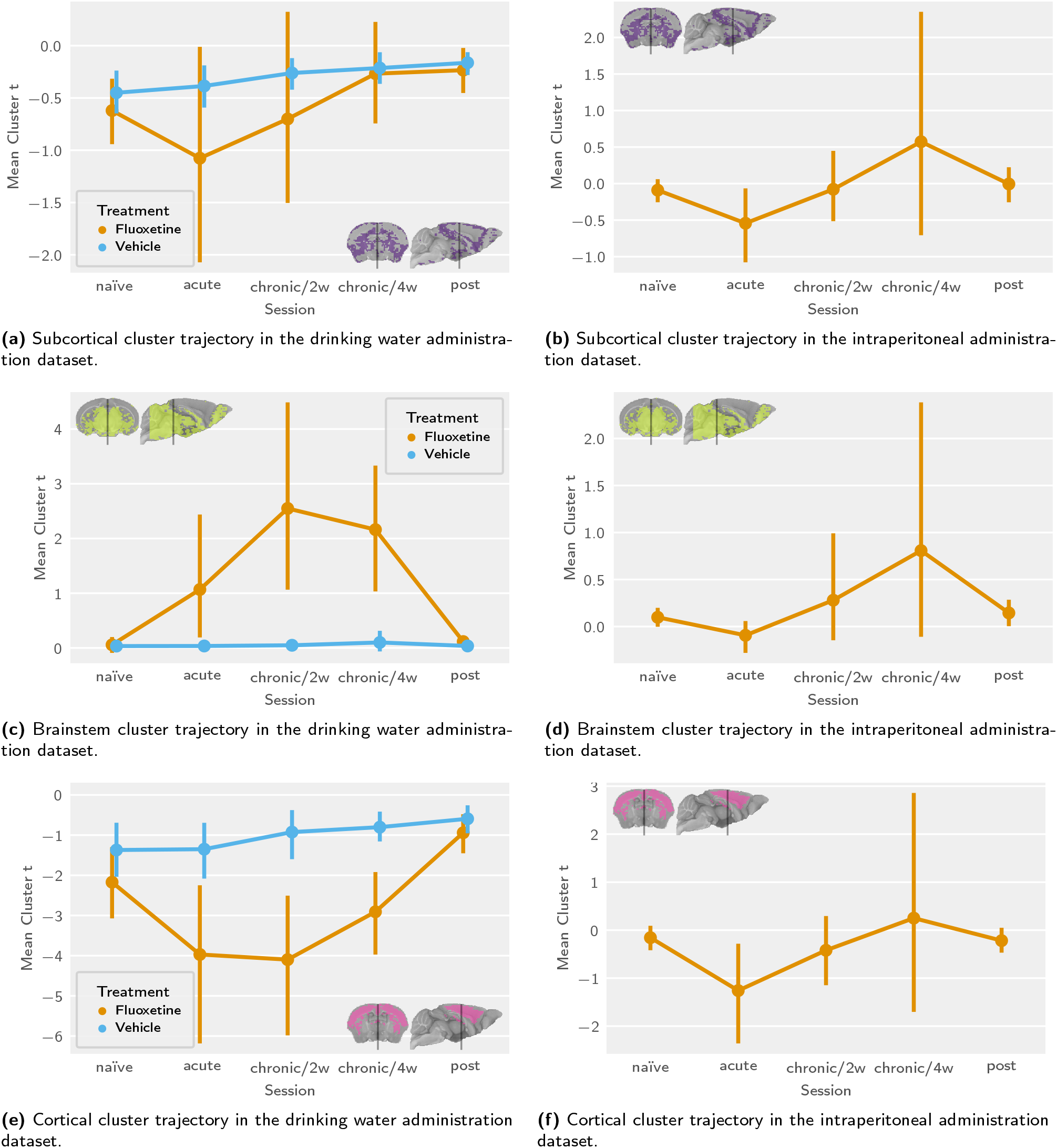
Seed-based analysis shows similar cluster distributions as stimulus-evoked analysis, with less distinct temporal profiles, and no discernible trajectory correspondence in the intraperitoneal administration data. Presented are longitudinal summaries of cluster t-statistic means from first level seed-based analysis, for both the drinking water and intraperitoneal administration datasets. The clusters used are classified based on of fluoxetine-treatment session-wise second-level analysis results, analogous to the classification for the stimulus-evoked analysis (fig. 3b).

